# TrieDedup: A fast trie-based deduplication algorithm to handle ambiguous bases in high-throughput sequencing

**DOI:** 10.1101/2022.02.20.481170

**Authors:** Jianqiao Hu, Sai Luo, Ming Tian, Adam Yongxin Ye

## Abstract

**Background:** High-throughput sequencing is a powerful tool that is extensively applied in biological studies. However, sequencers may produce low-quality bases, leading to ambiguous bases, ‘N’s. PCR duplicates introduced in library preparation should usually be removed in genomics studies, and several deduplication tools have been developed for this purpose. However, two identical reads may appear different due to ambiguous bases and the existing tools cannot address ‘N’s correctly or efficiently.

**Results:** Here we proposed and implemented TrieDedup, which uses trie (prefix tree) data structure to compare and store sequences. TrieDedup can handle ambiguous base ‘N’s, and efficiently deduplicate at the level of raw sequences. We also reduced its memory usage by approximately 20% by implementing restrictedDict in Python. We benchmarked the performance of the algorithm and showed that TrieDedup can deduplicate reads up to 270-fold faster than pairwise comparison at a cost of 32-fold higher memory usage.

**Conclusions:** TrieDedup algorithm may facilitate PCR deduplication, barcode or UMI assignment and repertoire diversity analysis of large scale high-throughput sequencing datasets with its ultra-fast algorithm that can account for ambiguous bases due to sequencing errors.

**Availability:** TrieDedup is available at https://github.com/lolrenceH/TrieDedup

## 1 Background

High-throughput sequencing methods have been adapted and applied in many fields of biological studies, including immune repertoire studies (Lin *et al*., 2016; Chen *et al*., 2020). Polymerase chain reaction (PCR), used during high-throughput sequencing library preparation, may lead to overrepresented templates, when multiple copies of the same DNA templates are amplified. These identical reads are termed as PCR duplicates (Li *et al*., 2009). PCR amplification may be biased based on the sequence and quantity of DNA templates; therefore, PCR duplicates usually need to be marked or removed to keep only one copy of their original template, through deduplication. However, high-throughput sequencing has a relatively higher sequencing error rate in comparison to traditional Sanger sequencing, which poses challenges for data analysis, including deduplication. High-throughput sequencers use base quality score (*Q* score) to represent their confidence in the identity of each base (Cock *et al*., 2010). *Q* scores are logarithmically related to the base calling error probabilities *P*, such that *Q* = -10×log_10_*P* (Ewing and Green, 1998). For example, a *Q* score of 10 represents an estimated sequencing error rate of 10%, and a *Q* score of 20 represents an error rate of 1%. Due to Illumina sequencing chemistry, the average base quality usually decreases from 5’-end to 3’-end of the reads (Manley *et al*., 2016). Low-quality bases, often considered as bases whose *Q* scores are below 10 or 20, can be conventionally converted to the ambiguous base ‘N’s (Hannon, 2010; Li, 2018). Low-quality reads with too many ‘N’s are often discarded as a means of quality control (Hannon, 2010). As described below, the presence of ambiguous ‘N’s in the remaining sequencing reads complicates the deduplication process.

Several bioinformatics tools have been developed for deduplication which work at two distinct levels, the level of alignment results and the level of raw reads. Tools that work on sequencing alignment files include samtools-rmdup (Li *et al*., 2009), Picard-MarkDuplciates (Broad Institute, 2019), EAGER-DeDup (Peltzer *et al*., 2016), and gencore (Chen *et al*., 2019). A common deduplication strategy of alignment-result-based tools is to drop the reads that have the same coordinates of read alignment, sometimes ignoring the underlying sequences or ambiguous bases, ‘N’s. Our previous bioinformatics pipeline for LAM-HTGTS also performs deduplication according to the alignment coordinates without considering the underlying sequences (Hu *et al*., 2016). On the other hand, tools that work at the level of raw sequencing data usually perform sequence comparisons and store the unique, deduplicated sequences in a hash data structure. Such tools include pRESTO (Vander Heiden *et al*., 2014), clumpify (Bushnell, 2021), and dedupe (Gregg and Eder, 2022). However, most sequence-based deduplication tools cannot handle ambiguous base ‘N’s correctly. In order to use hash for exact sequence matching, which is efficient when handling a large amount of data, they treat ‘N’ as a different base from regular bases ‘A’, ‘C’, ‘G’, ‘T’. Hence, sequence-based tools routinely consider two reads that only differ at positions of low sequencing quality as two distinct reads. The only one exception is the tool, pRESTO. pRESTO uses hash to store deduplicated sequences, and its implementation of a pairwise comparison algorithm can handle ambiguous base ‘N’s when comparing a query sequence to the stored deduplicated sequences one-by-one. However, pairwise comparison has a complexity of approximately O(n^2^), and may not be feasible due to the long processing time when dealing with large amounts of input sequences.

In B cell receptor repertoire studies, we are particularly interested in studying the diversity of V(D)J rearrangements and somatic hypermutations (Lin et al., 2016; Chen et al., 2020), where we need to distinguish unique sequences from potential PCR duplicates. To better investigate the B cell receptor repertoire, we have developed high-throughput genome-wide translocation sequencing-adapted repertoire and somatic hypermutation sequencing (HTGTS-Rep-SHM-Seq) assay, which can cover nearly full-length of the V(D)J-rearranged sequences after merging paired-end long-length MiSeq reads (Chen et al., 2020). This assay is based on the genomic DNA sequence in B cells. Each B cell has at most one productive V(D)J rearranged allele for heavy chain or light chain; therefore, after deduplication, each read of productive V(D)J rearrangement will represent one B cell. Due to the high error rate of next generation sequencing technology, we cannot avoid low-quality, ambiguous bases in the long reads. Low-quality reads with too many ‘N’s are often discarded as a means of quality control (Hannon, 2010). However, if we only keep the reads without any ‘N’s, rare events detected by a few reads with ‘N’s may also be discarded. Such rare events are often important in the repertoire diversity studies. We have observed otherwise identical reads that only differed at a few low-quality bases or ambiguous ‘N’s. They are likely to be duplicates of the same template, and, therefore, need to be deduplicated. For efficiency in processing huge amounts of sequencing data, our previous pipeline for HTGTS-Rep-SHM-Seq uses an alignment-result-based approach. It deals with ‘N’s by separating reads with and without ‘N’s, aligning reads with ‘N’s to reads without any ‘N’s using bowtie2, and checking their alignment length for deduplication (Chen et al., 2020). However, this approach cannot deduplicate among reads with ‘N’s when they do not have common equivalent reads without any ‘N’s. On the other hand, by pairwise comparison, pRESTO can deduplicate among reads with ‘N’s; but it runs slowly with the tremendous amount of input sequences.

Here, we designed and implemented TrieDedup, a faster deduplication algorithm that uses trie (prefix tree) structure to store deduplicated sequences and efficiently deduplicates at the level of raw sequences, ignoring differences only due to low-quality ambiguous base ‘N’s. We implemented a custom Python class, restrictedDict, to reduce memory usage. We benchmarked the performance of TrieDedup and the pairwise comparison algorithm implemented in pRESTO with simulated data as well as real public data. The source code of TrieDedup is available at https://github.com/lolrenceH/TrieDedup under Apache 2.0 license.

## 2 Implementation

### 2.1 Deduplication algorithm

Many sequence-based deduplication tools regard the ambiguous ‘N’ as different from the traditional bases, ‘A’, ‘C’, ‘G’, ‘T’, using hash-based exact matching to perform deduplication. Hash algorithm is highly efficient for comparing the literal identity of sequences. However, it offers no room for correctly accounting for sequencing ambiguity. Ambiguous ‘N’s potentially represent any of the four regular DNA bases. They should not be considered as different from other DNA bases by default.

Accounting for ‘N’s in deduplication poses two challenges: (1) when allowing differences at ‘N’s, the equivalence relationship between sequences may become complicated; and (2) we need an efficient algorithm to compare between a large amount of sequences and ignore ‘N’ differences.

For Challenge (1), theoretically, a network graph of equivalence relationship can be constructed, where each node represents an input sequence, and equivalent nodes are connected by an edge. Deduplication can be regarded as the well-known ‘maximal independent set (MIS)’ problem on the graph. A MIS is a set of nodes that are not adjacent, and its members and their neighbors include all the nodes in the graph. A deduplicated set is equivalent to a MIS on the network graph. The complication is that MIS may be not unique, and the sizes of MISs may vary. As a toy example, a simple equivalence graph ‘TAC’-’TNC’-’TGC’ has an MIS {‘TAC’,’TGC’} and another MIS {‘TNC’}. More generally, a star-shaped graph can have an MIS consisted of the tip nodes, or another MIS consisted of the center node. Thus, we need a principle for choosing of a MIS. For sequence deduplication, we may prefer to choose the nodes with fewer ‘N’s to represent the observed sequences, which correspond to the tip nodes of star-shaped graph.

Finding a MIS can be achieved by adding a candidate node into a MIS and removing neighbors of the node from the query, iteratively. Instead of performing pairwise comparison between all input sequences, we can store unique sequences that are previously deduplicated and compare each query sequence to these established deduplicated sequences, reducing the number of comparisons (Fig. 1). Because we prefer unambiguous sequences, we sort the input sequences by the number of ‘N’s in an ascending order, and consider each read, sequentially. This progressive pairwise comparison is implemented in pRESTO.

**Fig. 1.**
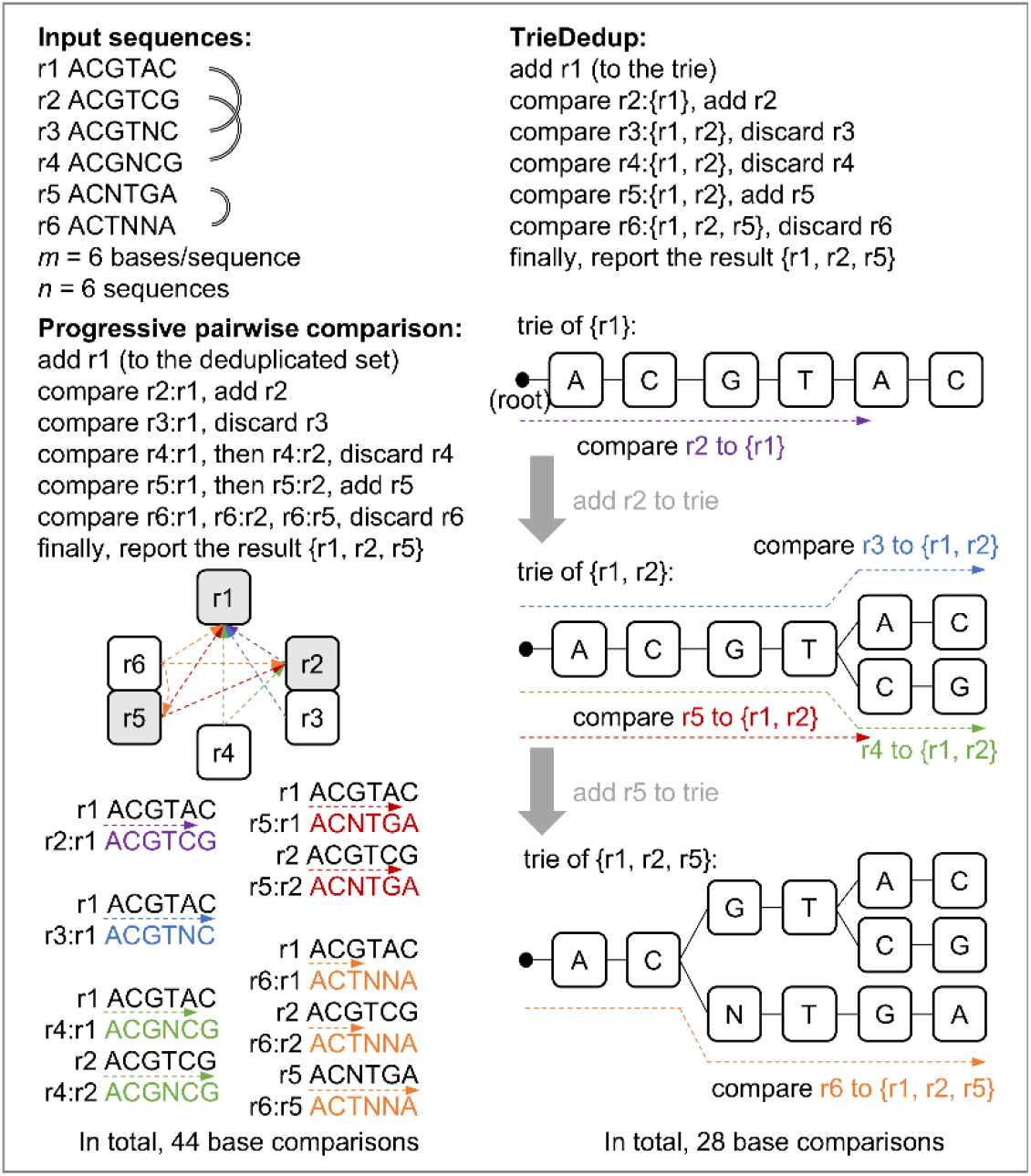
Diagram of progressive pairwise comparison and TrieDedup algorithm.

For Challenge (2), the pairwise comparison algorithm needs to compare each input sequence with the full set of deduplicated sequences to determine if it is unique. We adapted the trie (prefix tree) structure to store the previously deduplicated sequences, whose prefixes are organized into a consensus tree. The trie structure can retain information of sequence similarity from previous comparisons, thereby reducing the number of necessary comparisons. The trie structure can immediately identify an unobserved sequence, as soon as the input sequence diverges from the observed paths, thus reducing the number of comparisons (Fig. 1).

As a brief summary, we designed and implemented the following algorithms to store and compare sequences, which can ignore mismatches due to ‘N’s.

#### Algorithm 1.

Deduplication with trie storing a working set of deduplicated sequences

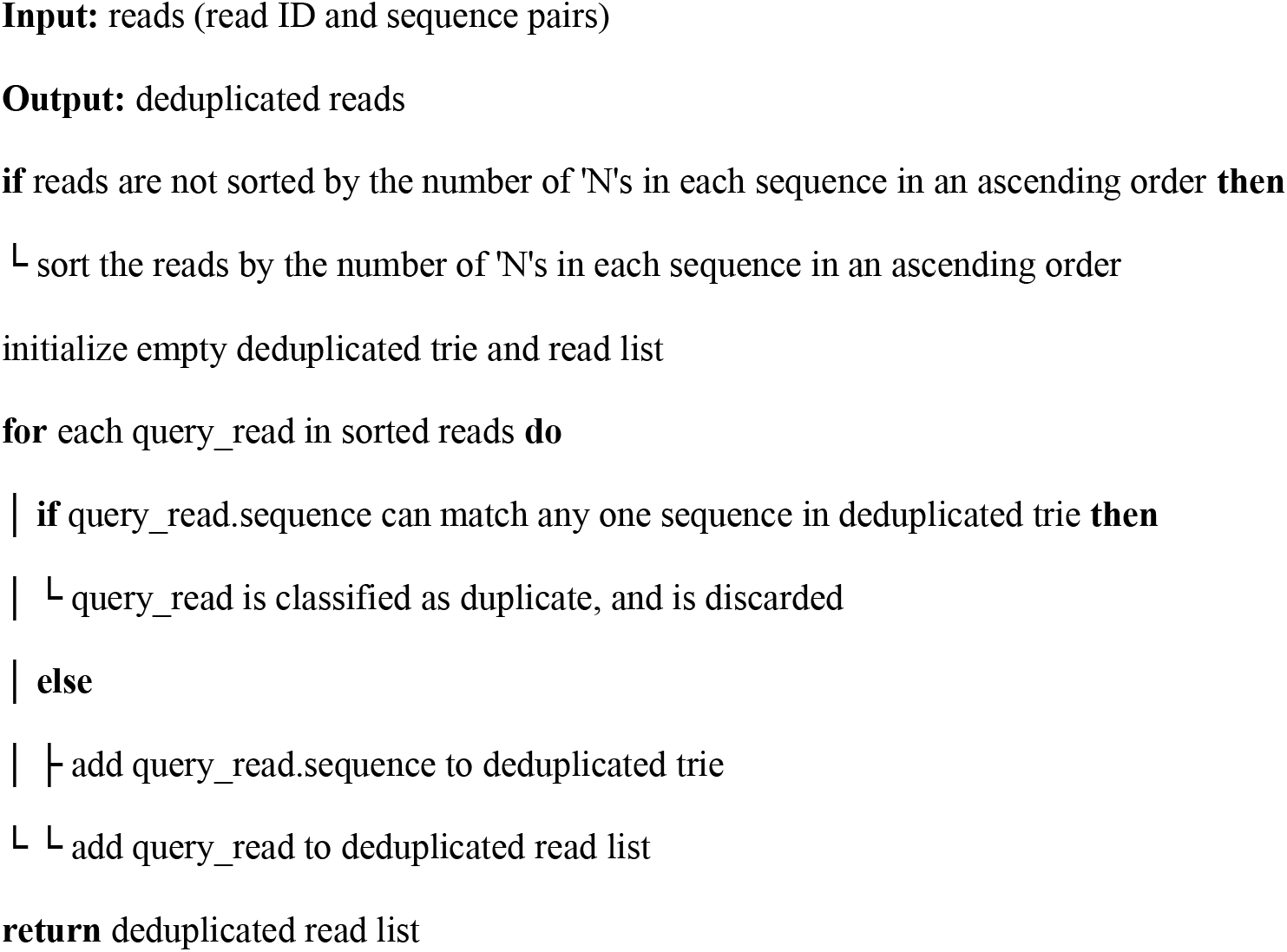

#### Algorithm 2.

Adding a sequence to the trie storing already deduplicated sequences

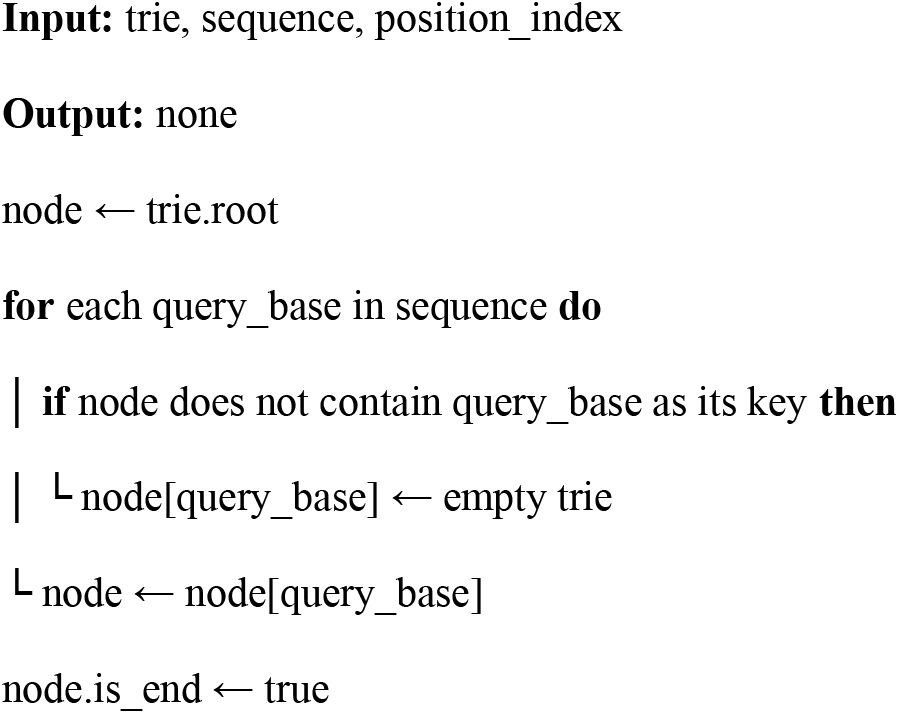

#### Algorithm 3.

Searching for a query sequence in the trie storing already deduplicated sequences

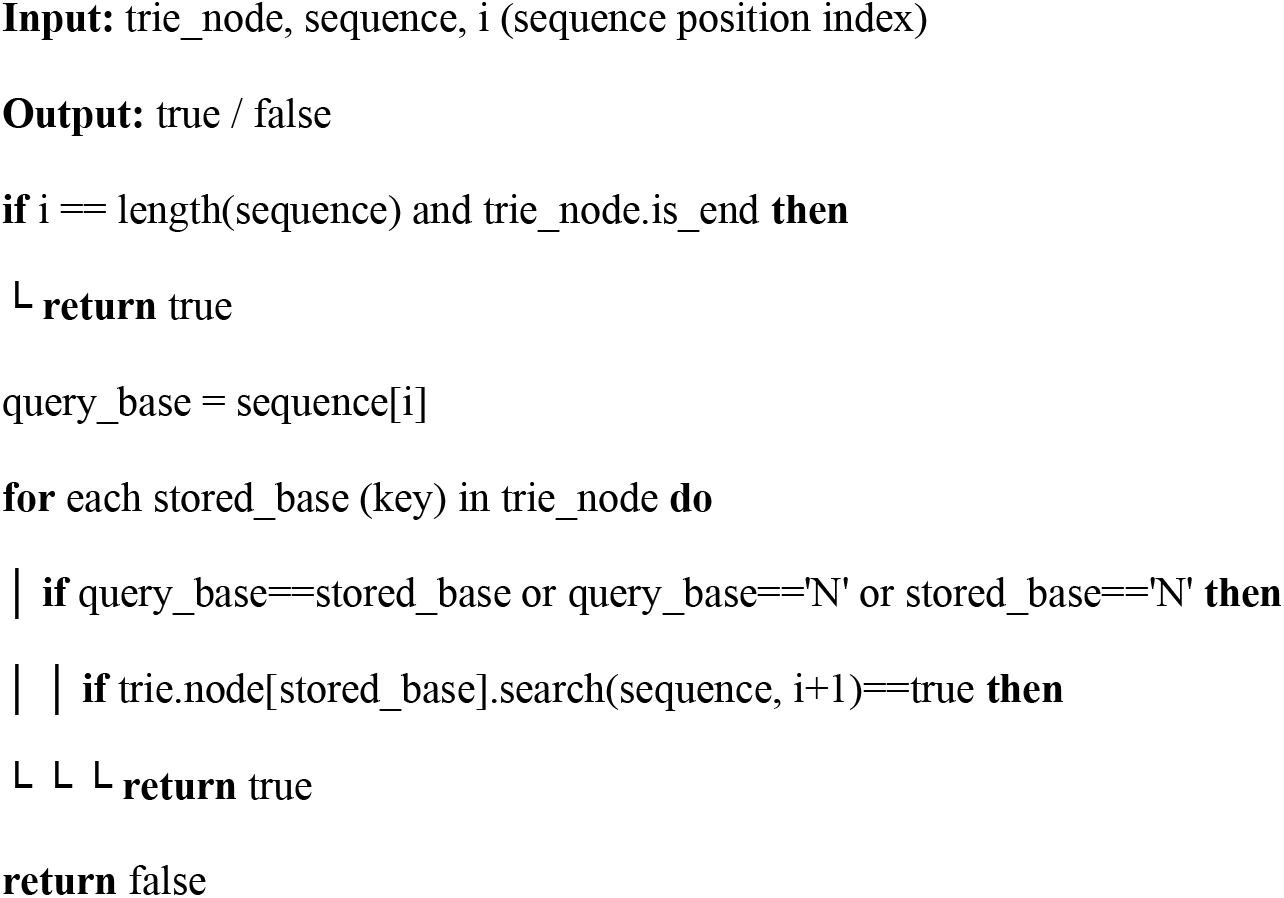

Note:

1. keep ‘N’ the last stored_base (after regular bases) when traversing trie_node
2. searching the whole sequence from trie_root by: trie_root.search(sequence, 0)

### 2.2 Memory usage optimization

The pairwise comparison algorithm directly stores every deduplicated, unique sequences, while our algorithm stores them in the trie structure, which theoretically costs less memory. However, empirically, storing a trie structure may cost more memory on the dict data structure than just storing simple sequences in hash keys, potentially due to Python’s base-level optimizations. To reduce memory consumption, we used__slots__magic in python to declare the variables, instead of storing them in another dict by default. We also implemented restrictedDict, a restricted dict with list implementation: it restricts hash keys to predetermined options, stores the values in a list instead of a dict, and keeps a shared dict to map limited keys to the index in the list. The restrictedDict achieves a smaller memory usage by storing values in a list instead of a dict, accommodating for large sequencing datasets. In DNA sequencing data, there are five possible base identities: ‘A’, ‘C’, ‘G’, ‘T’ and ‘N’. Thus, we only keep one dict as a shared class variable of restrictedDict, which stores the mapping table from the base letter (as the keys of dict) to the index in the list. For simplicity in describing the results, we abbreviate the original trie implementation as trie0, trie with only slots magic as trie1, and trie with slots and restrictedDict as trie2.

### 2.3 Benchmark and comparison to other tools

We compared progressive pairwise comparison algorithm and trie implementation with or without memory optimization in a Dell PowerEdge T640 server, with 192GB physical memory and limited to one CPU of Intel Xeon Gold 6126 2.6GHz, and in RedHat Linux operation system. We reimplemented the progressive pairwise comparison algorithm according to pRESTO. We benchmarked the running time and memory usage using both simulated reads or real public reads as input.

In order to simulate input sequences, we first generated a unique set of parental sequences of a specific length, then randomly sampled them with replacement, which usually sampled approximately 55% of unique parental sequences, and lastly converted a fixed number of bases at random locations to ‘N’s in each sampled sequence. We regarded the sequences with ‘N’s from the same parental sequence as PCR duplicates; therefore, we regarded the amount of unique parental sequences as the truth of the size of deduplicated set. We performed comparisons for 10^3^, 2×10^3^, 5×10^3^, 10^4^, 2×10^4^, 5×10^4^, 10^5^, 2×10^5^, 5×10^5^, and 10^6^ input sequences, with length of 30, 100, 150, and 200 bp, and converted 1%, 5%, 10% or 20% bases to ‘N’s for each input sequence. Each condition is tested with three repeats.

We also benchmarked the performance in public sequencing data. We downloaded SRR3744758, SRR3744760 and SRR3744762 raw fastq files from SRA, and randomly retrieved 10^6^ reads and masked bases whose quality score ≤ 10 by ‘N’ as the input sequences.

## 3 Results

### 3.1 Theoretical complexity analysis

Suppose there are *n* input sequences, and each sequence has *m* bases. For the preprocessing steps, the time complexity of counting ‘N’s is O(*m*×*n*), and sorting *n* sequences can be O(*n*×log(*n*)) for quick sort, or O(*n*) for bucket sort.

Suppose we progressively add sequences to the deduplicated set, and the deduplicated set has already stored *n*_d_ sequences. The space complexity of plain storing *n*_d_ sequences is O(*m*×*n*_d_). The algorithm of pairwise comparison between a candidate sequence and the deduplicated set has time complexity O(*m*×*n*_d_). Then, the overall complexity of the whole deduplication process is O(*m*×*n*^2^). This algorithm has been implemented in pRESTO, which runs slowly with a large number *n* of input sequences.

Here, we designed and implemented algorithm using the trie structure to store the already deduplicated sequences. The upper limit of the space complexity of trie structure storing *n*_d_ sequences is O(*m*×*n*_d_), and should be lower when the sequences have common prefixes. If there are no ‘N’s, the time complexity of comparison between a query sequence and the trie structure is only O(*m*); therefore, the lowest overall complexity of the whole deduplication process is O(*m*×*n*). However, when allowing ambiguous base ‘N’s, we may need to explore more branches in the trie to determine the comparison result. Theoretically, the upper limit of complexity for one query sequence is O(*m*×*n*_d_); therefore, the upper bound of the overall complexity is still O(*m*×*n*^2^). Though, this situation may rarely happen as long as the input sequences do not contain too many ‘N’s. If each sequence has at most *k* ‘N’s, the upper limit of the time complexity between one query sequence and the trie structure is O(*m*×5^*k*^), for 5 possible choices of bases (‘A’, ‘C’, ‘G’, ‘T’, ‘N’) at *k* trie nodes; therefore, overall time complexity is O(*m*×*n*×5^*k*^). Theoretically, the actual time complexity is dependent on the amount and location of ‘N’s in sequences. Fewer ‘N’s or ‘N’ locating closer to the end of sequences (near trie tips) will have less complexity than more ‘N’s or ‘N’ locating closer to the start of sequences (near trie root).

### 3.2 Running time and memory usage in simulated data

We benchmarked the speed and memory consumption of the trie algorithm with or without memory optimization, and compared them to the performance of the progressive pairwise comparison algorithm, which we reimplemented from pRESTO, using the simulated 200-bp reads. All the algorithms show an approximately linear relationship between log-transformed running time and the log-transformed number of input sequences (Fig. 2A). The slope of pairwise comparison is close to the theoretical order 2. On the other hand, the slope of trie algorithm is much lower, which is ≤ 1.3 for n ≤ 10^5^ and the percentage of ambiguous bases in reads (N%) ≤ 10%, and increases for larger *n* or N%. When the input sequences are less than 5000, the pairwise comparison algorithm is more efficient than the trie algorithm whose performance is less than 3s. When there are more than 5000 input sequences, the trie algorithm runs significantly faster than pairwise comparison.

**Fig. 2.**
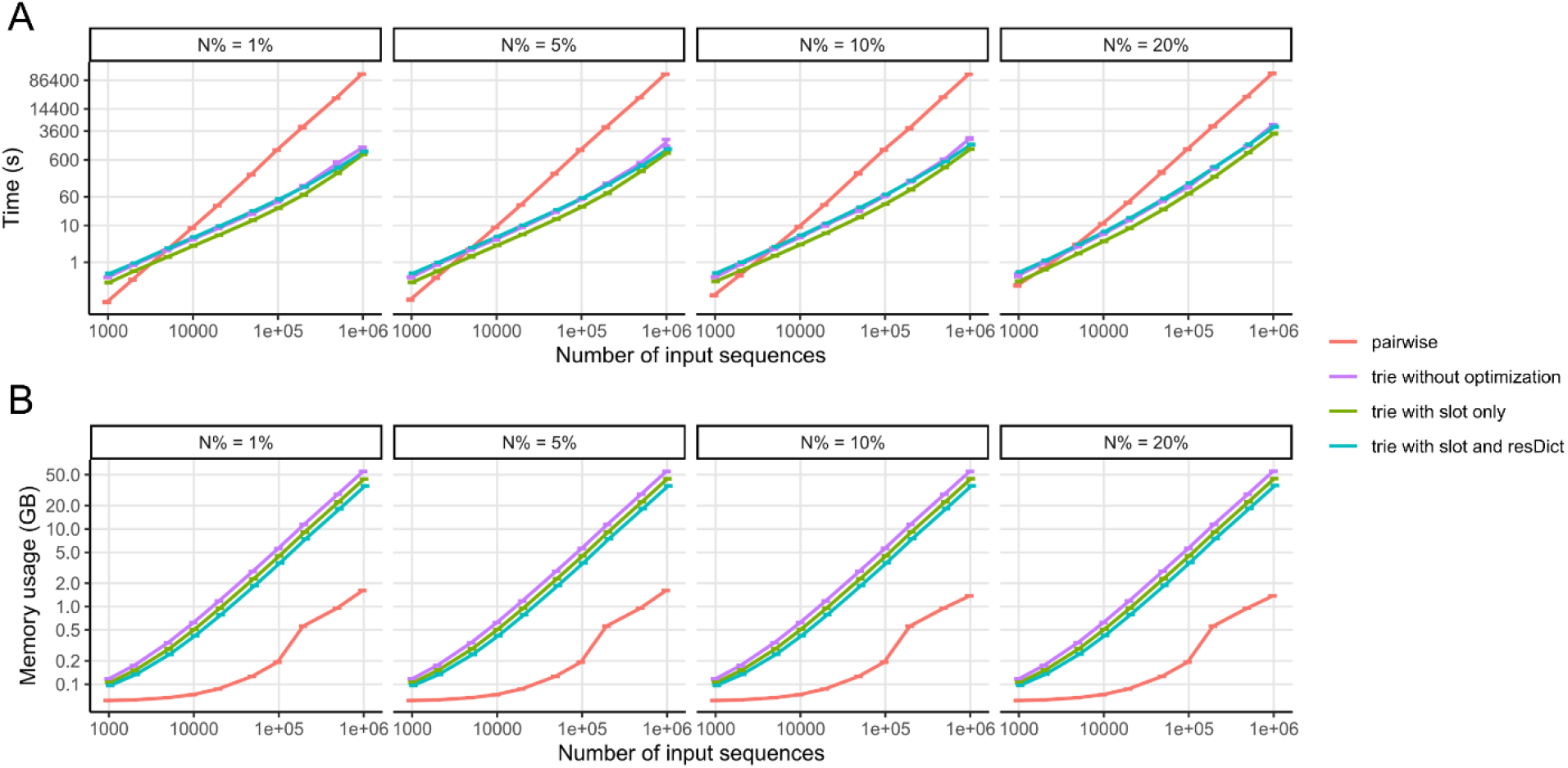
Running time and memory usage increases with larger amount of input sequences (benchmark simulation). (A) running time; (B) memory usage. Input sequences are 200 bp in length. Error bars shows mean ±standard deviation, each with 3 replicates.

The trie algorithm significantly outperforms pairwise comparison at large input sizes. For example, at N% = 5% and *n* = 10^4^, pairwise comparison needs 8.9s (on average), trie algorithm needs ≤ 5.0s; at *n* = 10^5^, pairwise comparison needs 1100 s (or 18.3 min), trie algorithm needs ≤ 55s; at *n* = 10^6^, pairwise comparison needs 125000s (or 34.7h), trie algorithm needs ≤ 1755 s (or 29.3 min).

When comparing the running time among the three different options of memory optimization of the trie implementation, we found that all the three options have very similar running time, without magnitude difference. Trie1 runs relatively faster than trie0 and trie2, with the difference being less than 2-fold (Fig. 2A).

On the other hand, from the perspective of memory usage, the pairwise algorithm is more memory-efficient to implement than the trie because entire sequences can be stored as one item instead of storing each base individually (Fig. 2B). Among three trie implementations, as expected, trie0 requires the most memory and trie2 requires the least. The slopes for the three options in Python implementation are similar, all close to but a little less than the theoretical 1. For *n* ≥ 10^5^, the implementation of restrictedDict (trie2) improves the memory usage of__slots__optimization (trie1) by approximately 19%, while trie1 improves 20% compared to trie0; therefore, trie2 only uses 65% memory as much as trie0 uses.

When N% = 5% and *n* = 10^4^, pairwise comparison on average requires 0.07GB memory to run, while trie0 requires 0.62GB, trie1 0.51GB, and trie2 0.42GB; when *n* = 10^5^, pairwise comparison needs 0.19GB, trie0 5.6GB, trie1 4.5GB, and trie2 3.7GB; when *n* = 10^6^, pairwise comparison requires 1.6GB, trie0 55GB, trie1 44GB, and trie2 36GB.

We also evaluated the influence of the length of input sequences and N% on the performance of pairwise comparison and trie2 (data not shown). As expected, longer length will need more running time and memory usage. The percentage of ambiguous bases in reads (N%) hardly affects the memory usage of both algorithms, nor does it influence the running time of pairwise comparison.

### 3.3 Running time and memory usage in real data

We applied the two algorithms to published sequencing data with a read length of 300 bp and an average N% of 4.9∼15.8%. As expected, higher N% and lower percentage of unique reads were observed in R2 than R1 reads. With an input size of 10^6^ raw reads, both pairwise comparison and trie algorithms reported the same number of unique reads (8.12∼9.48×10^5^). Deduplication by trie2 only needed 0.9∼2.1 hours using 35∼55GB of memory, while deduplication by pairwise comparison required 6∼16 days with about 1.5GB memory usage (Fig. 3). Therefore, trie2 deduplication can achieve about 270-fold faster speed than pairwise comparison, with 32-fold higher memory usage.

**Fig. 3.**
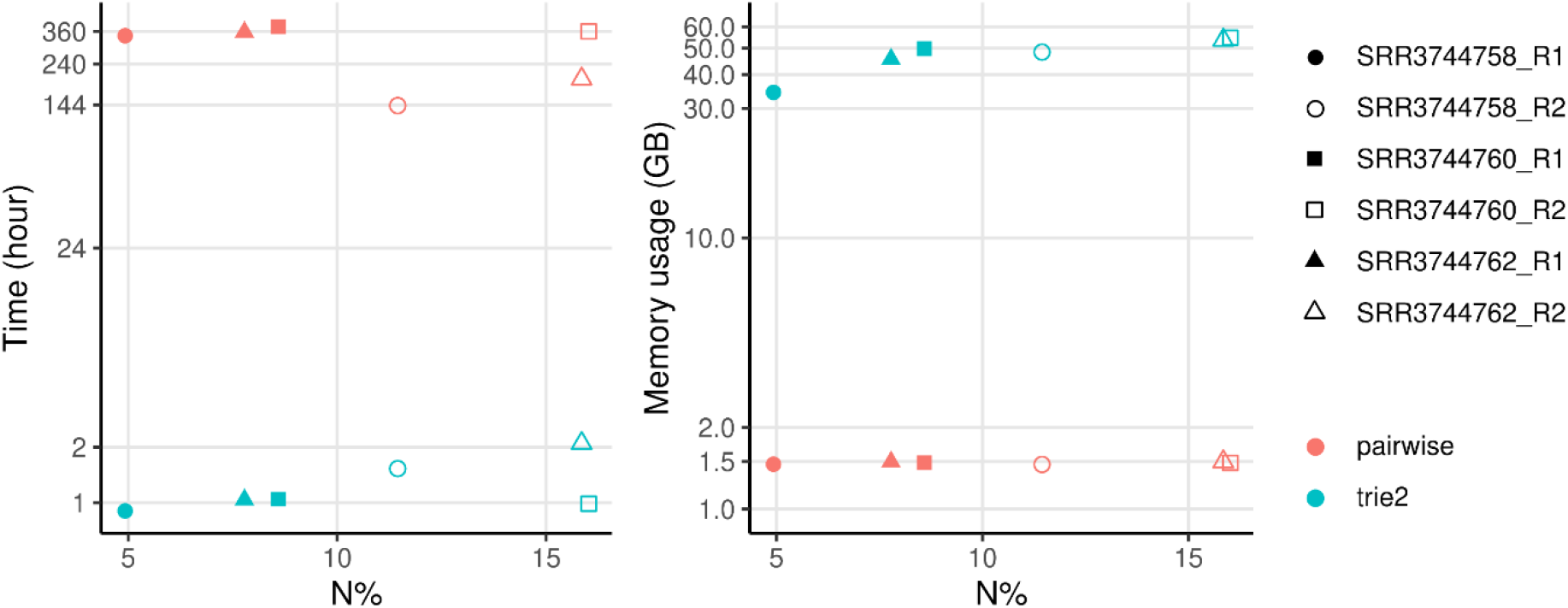
Running time and memory usage when applying on 10^6^ real 300-bp reads from SRA. X-axis ‘N%’ shows the average percentage of ambiguous base ‘N’s in reads

## 4 Discussion

To deduplicate high-throughput sequencing libraries while ignoring differences only due to ambiguous base ‘N’s, we adapted trie structure to store deduplicated sequences, and implemented corresponding algorithm, named as TrieDedup. When the input size is larger than 5000 sequences, the trie algorithm is more efficient than the pairwise comparison algorithm used in pRESTO, at the price of higher memory usage. TrieDedup can deduplicate up to 10^6^ input sequences within 2 hours using less than 36GB of memory. In addition, TrieDedup may be potentially adapted into pRESTO framework.

Because the running time and memory usage both increased with more input sequences, and for running time the complexity order on N is larger than one, the efficiency of deduplication may be further improved if we can divide the input sequences into smaller non-overlapping groups. For example, the reads may be divided into groups based on the coordinate if the reads have been aligned, or the V and J alignment for V(D)J repertoire. If these subsets of data are processed separately, the algorithm will have to store fewer reads simultaneously, hence reducing memory usage.

Trie structure has also been used in deduplication of Unique Molecular Identifiers (UMIs) (Liu, 2019), but the traditional trie structure cannot handle ambiguous bases, although errors in UMIs are common (Smith *et al*., 2017). UMIs containing any ‘N’s or bases with *Q* score < 10 are by default filtered out in 10x Genomics Cell Range processing. Here, we designed and implemented TrieDedup, where the specialized trie structure and algorithm can correctly and efficiently handle the differences due to ambiguous bases. With its ultra-fast algorithm, TrieDedup may also potentially be applied to barcode or UMI assignment when considering reads with a few low-quality bases in the UMIs.

Suffix tree may also be used instead of trie (prefix tree) with similar algorithms. Because of the Illumina sequencing chemistry, sequencers tend to produce more ambiguous base ‘N’s towards the 3’-end of sequences (trie tips), rather than the 5’-end (trie root). Therefore, we chose trie which theoretically have less complexity than suffix tree. In our benchmark, we noticed both pairwise comparison and TrieDedup perform faster when ‘N’s are distributed near the 3’-end of a read, in comparison to being evenly distributed across a read. The improvement of speed is larger for TrieDedup than pairwise comparison (data not shown). In the real data of 300-bp reads, most ‘N’s are distributed in the 100-bp part near the 3’-end, and TrieDedup run faster than pairwise comparison for even just 1000 input sequences.

We did not restrict the allowed keys of restrictedDict to ‘A’, ‘C’, ‘G’, ‘T’ and ‘N’, for possible generalizability to other scenarios, such as amino acid sequences. To further improve on runtime and memory usage, we also implemented TrieDedup in Java and C++, and restricted the allowed keys to DNA bases. These implementations are also available in our GitHub repository. Compared to Python implementation, TrieDedup C++ implementation achieved 5∼11-fold faster and 1/3 memory.

For the comparison algorithm, other than the recursive version shown in Materials and methods, we have also designed, implemented and tested several iterative versions. However, those iterative versions have similar or poorer speed and memory performance (data not shown), and are hence omitted.

The threshold of *Q* score for converting low-quality bases to ambiguous ‘N’s, which is often library-specifically set to 10 or 20 arbitrarily, may affect N% in input reads, as well as the amount of deduplicated reads. A potentially more principled approach is to sum up the error rate of mismatches in pairwise comparison, and then set the threshold on the sum error rate to judge the equivalence between reads. But it may generate more complicated relationship of equivalence and even higher computational complexity than the current pairwise comparison algorithm in pRESTO.

## 5 Conclusions

We implemented TrieDedup, which uses trie structure to store deduplicated sequences, and ignores differences only due to ambiguous base ‘N’s. We also implemented a memory-efficient class, restrictedDict, that reduced the memory usage to about 0.8-fold. TrieDedup significantly outperforms the pairwise comparison strategy when the amount of input sequences is larger than a few thousand. TrieDedup can deduplicate reads up to 270-fold faster than pairwise comparison at a cost of 32-fold higher memory usage. Potentially, TrieDedup may be adapted into pRESTO, and may be generalized to other scenarios for deduplication with ambiguous letters.

## Availability and requirements

**Project name:** TrieDedup: A fast trie-based deduplication algorithm to handle ambiguous bases in high-throughput sequencing

**Project home page:** https://github.com/lolrenceH/TrieDedup

**Operating system(s):** Platform independent

**Programming language:** Python, with C++ and Java implementations available on GitHub

**License:** Apache 2.0

## List of abbreviations

PCR: Polymerase chain reaction
*Q* score: Base quality score
HTGTS-Rep-SHM-Seq: High-throughput genome-wide translocation sequencing-adapted repertoire and somatic hypermutation sequencing
MIS: Maximal independent set
trie0: trie implementation without memory optimization
trie1: trie implementation with only__slots__magic
trie2: trie implementation with__slots__and restrictedDict
N%: Percentage of ambiguous bases in reads
UMIs: Unique Molecular Identifiers

## Declarations

### Ethics approval and consent to participate

Not applicable.

### Consent for publication

Not applicable

### Availability of data and materials

TrieDedup code is available at https://github.com/lolrenceH/TrieDedup

### Authors’ contributions

SL and MT raised the scientific questions, AYY conceptualized and designed the software. AYY and JH developed the software. JH, SL and MT, and AYY tested and validated the software. JH performed benchmark analysis. JH and AYY conducted data analysis and visualization. JH, SL and MT, and AYY wrote the manuscript. All authors read, critically revised, and approved the final manuscript.

## Acknowledgements

We would like to express our deep gratitude to Prof. Frederick W Alt, for his unwavering support and generously providing the necessary resources and funding throughout this projects.

## Open access

For the purpose of open access, the author has applied a CC BY 4.0 public copyright license to any author accepted manuscript version arising from this submission.

## References

Broad Institute (2019) Picard toolkit. https://broadinstitute.github.io/picard/.

Bushnell, B. (2021) BBMap - clumpify. https://github.com/BioInfoTools/BBMap/blob/master/sh/clumpify.sh.

Chen, H. et al. (2020) BCR selection and affinity maturation in Peyer’s patch germinal centres. Nature, 582, 421–425. doi:10.1038/s41586-020-2262-4.

Chen, S. et al. (2019) Gencore: an efficient tool to generate consensus reads for error suppressing and duplicate removing of NGS data. BMC Bioinformatics, 20, 606. doi:10.1186/s12859-019-3280-9.

Cock, P.J.A. et al. (2010) The Sanger FASTQ file format for sequences with quality scores, and the Solexa/Illumina FASTQ variants. Nucleic Acids Research, 38, 1767–1771. doi:10.1093/nar/gkp1137.

Ewing, B. and Green, P. (1998) Base-Calling of Automated Sequencer Traces Using Phred. II. Error Probabilities. Genome Res., 8, 186–194. doi:10.1101/gr.8.3.186.

Gregg, F. and Eder, D. (2022) Dedupe. https://github.com/dedupeio/dedupe.

Hannon, G.J. (2010) FASTX-Toolkit. http://hannonlab.cshl.edu/fastx_toolkit.

Hu, J. et al. (2016) Detecting DNA double-stranded breaks in mammalian genomes by linear amplification– mediated high-throughput genome-wide translocation sequencing. Nature Protocols, 11, 853–871. doi:10.1038/nprot.2016.043.

Li, H. (2018) seqtk. https://github.com/lh3/seqtk.

Li, H. et al. (2009) The Sequence Alignment/Map format and SAMtools. Bioinformatics, 25, 2078–2079. doi:10.1093/bioinformatics/btp352.

Lin, S.G. et al. (2016) Highly sensitive and unbiased approach for elucidating antibody repertoires. Proceedings of the National Academy of Sciences, 113, 7846–7851. doi:10.1073/pnas.1608649113.

Liu, D. (2019) Algorithms for efficiently collapsing reads with Unique Molecular Identifiers. PeerJ, 7, e8275. doi:10.7717/peerj.8275.

Manley, L.J. et al. (2016) Monitoring Error Rates In Illumina Sequencing. J Biomol Tech, 27, 125–128. doi:10.7171/jbt.16-2704-002.

Peltzer, A. et al. (2016) EAGER: efficient ancient genome reconstruction. Genome Biol, 17, 60. doi:10.1186/s13059-016-0918-z.

Smith, T. et al. (2017) UMI-tools: modeling sequencing errors in Unique Molecular Identifiers to improve quantification accuracy. Genome Res., 27, 491–499. doi:10.1101/gr.209601.116.

Vander Heiden, J.A. et al. (2014) pRESTO: a toolkit for processing high-throughput sequencing raw reads of lymphocyte receptor repertoires. Bioinformatics, 30, 1930–1932. doi:10.1093/bioinformatics/btu138.

